# SSIM can robustly identify changes in 3D genome conformation maps

**DOI:** 10.1101/2021.10.18.464422

**Authors:** Elizabeth Ing-Simmons, Nick Machnik, Juan M. Vaquerizas

## Abstract

We previously presented Comparison of Hi-C Experiments using Structural Similarity (CHESS), an approach that applies the concept of the structural similarity index (SSIM) to Hi-C matrices^1^, and demonstrated that it could be used to identify both regions with similar 3D chromatin conformation across species, and regions with different chromatin conformation in different conditions. In contrast to the claim of Lee et al.^2^ that the SSIM output of CHESS is ‘independent’ of the input data, here we confirm that SSIM depends on both local and global properties of the input Hi-C matrices. We provide two approaches for using CHESS to highlight regions of differential genome organisation for further investigation, and expanded guidelines for choosing appropriate parameters and controls for these analyses.

## Main text

Lee et al.^2^ applied CHESS to Hi-C data from a DLBCL patient and healthy control^3^, and compared this to results obtained by mixing these datasets in equal parts to create two shuffled datasets. They show that the SSIM profiles obtained for these comparisons are very similar. Since both patient and control datasets are derived from B cells, we expect genome organisation to be largely similar between them, with differences reflecting disease-specific changes. Therefore, while the shuffled datasets are almost identical, we also expect the patient-control comparison to have similar structures at most loci. We repeated the dataset mixing analysis in order to test this hypothesis (see Methods). In line with our expectations, while the resulting SSIM profiles are highly correlated (Pearson’s R = 0.98), the comparison of shuffled datasets has overall higher SSIM values (Figure 1A), and importantly, the distributions of SSIM values are significantly different (Wilcoxon signed-rank test p-value < 2.2 × 10^−16^), indicating a higher similarity between the shuffled datasets.

**Figure 1.**
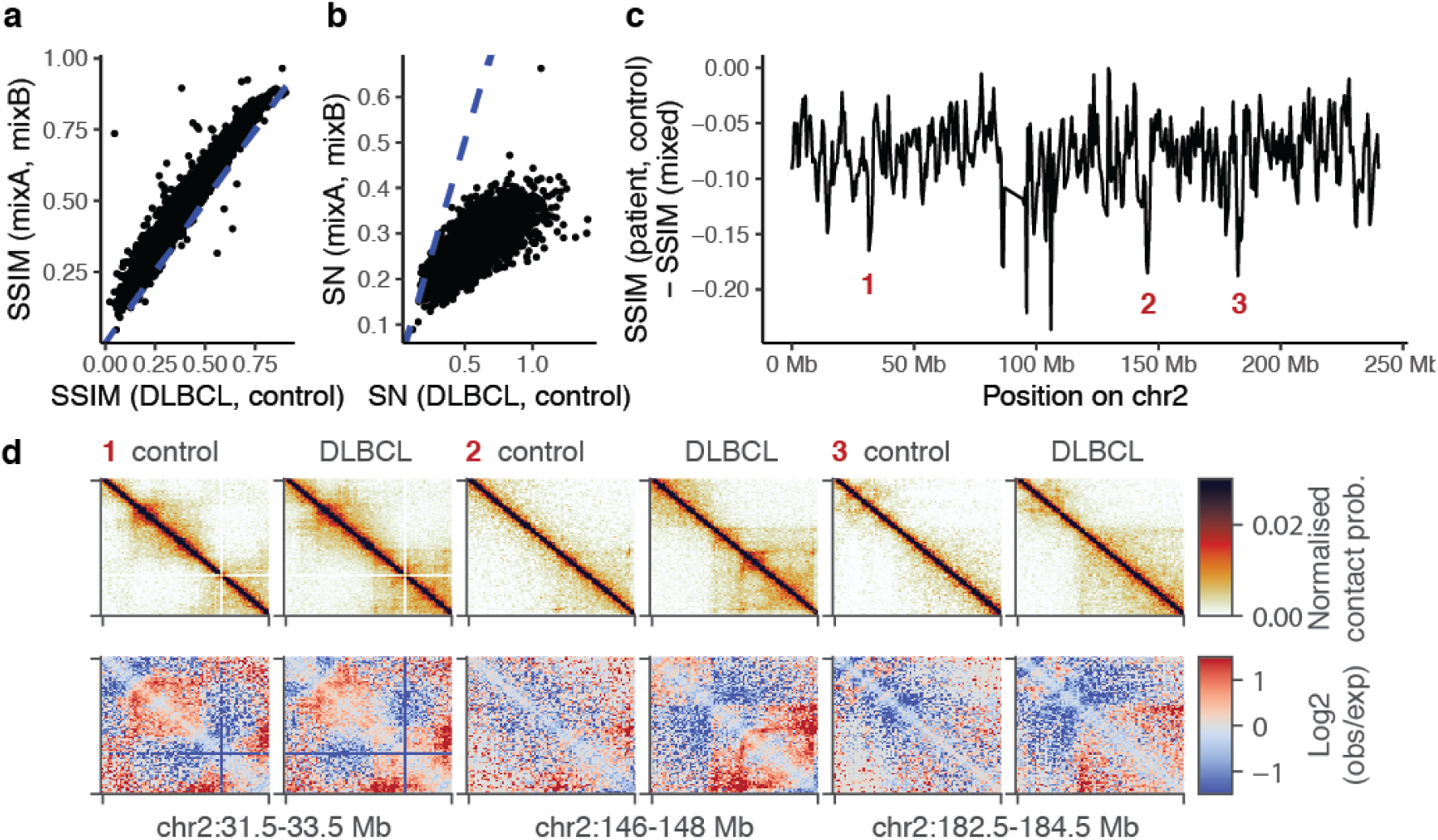
Differences in SSIM between the DLBCL-control comparison and the comparison of mixed datasets highlight changes in genome organisation. A. Scatterplot of the SSIM profile of the real DLBCL-control comparison versus the SSIM profile of the mixed comparison. B. Scatterplot of SN profile of the real DLBCL-control comparison versus the SN profile of the mixed comparison C. Subtraction of SSIM (mixed) from SSIM (DLBCL, control) highlights regions with changes between DLBCL and control. D. Example regions from C where the SSIM difference profile indicates differences between DLBCL and control. Top, normalised Hi-C data at 25kb resolution. Bottom, log2 observed / expected values for the same regions.

We also assessed the impact of noise in these comparisons. CHESS defines a signal-to-noise metric (SN), which measures the mean magnitude of values in a difference matrix of the input Hi-C datasets, divided by the variance across the matrix. We introduced this measure in the original CHESS implementation^1^ in order to control for noise effects, based on the observation that in noisy regions, values of neighboring pixels of a difference matrix often have opposite signs, fluctuating randomly, which leads to high variance. In regions with visually striking differences such as differential domain organisation, the values in the differential region of the matrix have low variance and high mean absolute value. The SN profiles of the patient-control comparison and the comparison of mixed datasets are correlated (Pearson’s R = 0.77), but the SN is lower in the comparison of mixed datasets (Wilcoxon signed-rank test p-value < 2.2 × 10^−16^), indicating that differences between the mixed datasets are largely due to noise (Figure 1B).

We next investigated the underlying cause of the correlations of SSIM and SN across different comparisons and the fluctuation of SSIM across the genome even for comparisons between theoretically highly similar datasets. We found that SSIM and SN correlate with both local Hi-C read depth and the variance of the insulation score in the window used for CHESS comparisons (Extended Data Figure 1A), although SSIM has a stronger correlation with insulation score variance than coverage. This suggests that genomic regions with low insulation score variance, indicative of a lack of domain structure, and lower Hi-C coverage, are more likely to have low SSIM, likely due to higher noise in these regions and lack of structure. These results are in agreement with our original examination of the role of noise in identifying similarities in Hi-C matrices^1^, and highlight the need to apply an SN threshold to identify regions with meaningful differences in the Hi-C data.

In addition, we assessed the relationship between total sequencing depth and SSIM and SN, by applying CHESS to comparisons of biological replicates with different sequencing depths (Extended Data Figure 1B-C). Comparisons involving replicates with higher sequencing depth have higher overall SSIM scores and lower variation in SN. This is likely due to lower levels of noise in these datasets. Therefore, care should be taken when comparing SSIM scores from comparisons involving datasets with unequal sequencing depths.

While there is fluctuation of the SSIM profile across the genome due to local effects, we reasoned that taking the difference between the SSIM profile of the DLBCL-control comparison and the SSIM profile of the comparison of mixtures would control for these local influences and identify regions with biological relevant reductions in SSIM in the real comparison (Figure 1C). Indeed, the local minima highlighted in Fig 1C correspond to regions with visible differences in chromatin conformation (Figure 1D, Extended Data Figure 2). This analysis is similar to that carried out in our previous publications ^1,4^ where we used the SSIM profile of a comparison between two control samples as a reference to identify regions with changes in chromatin conformation in an experimental condition compared to a control. This analysis confirms that the SSIM output of CHESS can be used to identify regions with differences in chromatin conformation.

Lee et al. also report that the SSIM profile is also highly correlated between Hi-C datasets produced from different cell lines^2^. Similarly to the DLBCL vs control datasets above, this is not surprising, since it has been shown before that domains and loops are conserved across human cell lines ^5,6^. However, as further validation that the SSIM profile produced by CHESS reflects differences in chromatin organisation, we carried out comparisons of Hi-C data from cohesin-depleted and CTCF-depleted cells with control cells^7^. Removal of cohesin or CTCF severely affects normal domain structure^8^. In addition, we compared data from cells in G1 with prometaphase, where chromatin is highly compacted and normal interphase structures are absent. As expected for such global perturbations, the mean SSIM values of these comparisons are markedly reduced and the correlation of SSIM with local Hi-C coverage was reduced or abolished (Extended Data Figure 3).

While taking the difference of SSIM profiles between a reference comparison and the comparison of interest highlights potential regions of interest, this approach does not make use of the SN value to remove regions where differences are due to noise. In our previous analysis of the DLBCL and control B cell data^1,3^ we applied an empirically determined SN threshold of 0.6, along with a SSIM Z-score threshold of -1.2, to select a subset of regions with high-confidence changes in genome organisation. However, since the global distributions of SSIM and SN depend on the sequencing depth of the samples (shown above), as well as on Hi-C matrix resolution and the window size used for CHESS (not shown), these thresholds are not universally applicable. Hi-C analysis approaches such as identification of TADs and loops often require investigators to choose appropriate thresholds or parameters which may be based on visual inspection of the data and algorithmic output ^9–11^; similarly, we advise users to choose CHESS parameters and SSIM and SN thresholds based on their data and the aims of their analysis.

Lee et al.^2^ used CHESS to compare publicly available Hi-C data from GM12878 and K562 cells using the same thresholds that we have used for the DLBCL and control B cell data above (SSIM Z-score < -1.2 and SN > 0.6). A quick examination of the distribution of SN values for the GM12878 and K562 comparison reveals that the choice of these thresholds is not suitable for these data, since the majority of the genome displays significantly higher SN values (Figure 2A), effectively resulting in a lack of filtering for noise in this analysis, and the subsequent artifactual identification of regions with dubious changes in 3D genome organisation.

**Figure 2.**
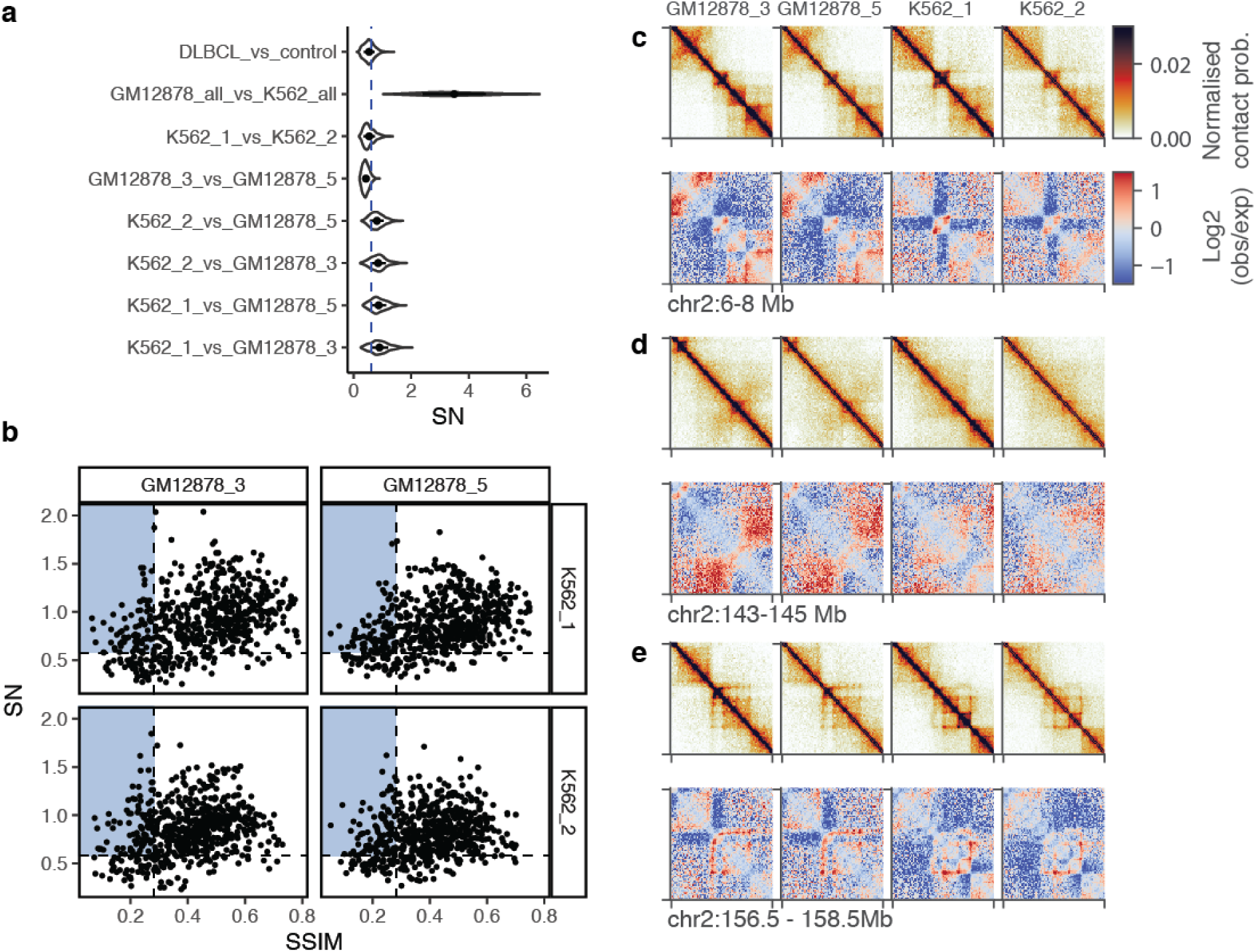
Data-based selection of SSIM and SN thresholds allows identification of windows with striking changes in genome organisation. A. SN distributions from various comparisons, demonstrating that these are highly variable. The dashed line indicates SN = 0.6, the threshold used in the original DLBCL-control comparison and used by Lee et al.^2^ for the comparison of GM12878 and K562 merged datasets. B. Scatterplot of SSIM against SN for comparisons between GM12878 and K562 individual biological replicates. Comparisons were carried out using Hi-C data downsampled to equal numbers of valid pairs, binned at 25 kb resolution, using window sizes of 2 Mb and a step size of 500 kb. Dashed lines indicate the 10th percentile of SSIM and the 90th percentile of SN derived from equivalent comparisons of biological replicates from the same cell line. Points in the blue shaded area were selected for further investigation. C-E. Example of regions from B selected for further investigation. Top, normalised Hi-C data at 25kb resolution. Bottom, log2 observed / expected values for the same regions.

To help with the selection of appropriate SN and SSIM thresholds, we sought to identify an approach that would produce adequate and interpretable data-based thresholds for SSIM and SN that do not solely rely on visual inspection of results. Differences between biological replicate Hi-C experiments should be minimal and due to noise. Therefore, we reasoned that the distributions of SSIM and SN produced by applying CHESS to biological replicates could be used to identify thresholds that would indicate output values that are unlikely to occur due to random variation between datasets. We compared two biological replicates of Hi-C from GM12878 and K562 cells, all downsampled to equal sequencing depth. We identified the 10th percentile of the SSIM distribution from these comparisons (GM12878: 0.274; K562: 0.282) and the 90th percentile of the SN distribution (GM12878: 0.577; K562: 0.813). Here we used the less stringent value of each pair as thresholds to identify regions that have lower SSIM and higher SN in comparisons between GM12878 and K562 than 90% of regions in comparisons of biological replicates of the same cell type (Figure 2B). None of the regions highlighted by Lee et al.^2^ pass these thresholds. Instead, 52 windows (out of 580 total 2 Mb windows on the chromosomes analysed here) passed the thresholds in all four pairwise comparisons; three of the top ten regions with lowest SSIM values are visualised in Figure 2C-E. These regions contain striking changes in domains, compartments, and loops, demonstrating the ability of CHESS to identify meaningful changes in 3D genome organisation and to filter regions dominated by noise, when appropriate thresholds are used. We note that this heuristic approach may not be suitable for all datasets, and investigators should carefully consider their individual threshold choices.

Since the development of Hi-C and the first algorithms for identification of domains, loops, and compartments, increasingly sophisticated approaches for detecting these features have been developed^12^. We expect that in the coming years further methods will be developed for the unbiased assessment of chromatin conformation without the necessity of first defining a structure of interest. As a method which analyses broad windows we believe CHESS is useful as a complement to unbiased methods that assess pixel-wise differences in Hi-C matrices, such as diffHiC^13^ or ACCOST^14^. Further analyses will increase understanding of the relationship between SSIM, SN, and chromatin conformation differences.

Overall, as opposed to the results reported by Lee et al.^2^, our results demonstrate that, when used appropriately, CHESS and SSIM can be employed to robustly identify changes in three-dimensional chromatin conformation.

## Methods

### Hi-C analysis

DLBCL patient and control Hi-C datasets were obtained from Diaz et al. 2018^3^ and processed as previously described^4^ using FAN-C^15^ version 0.9.18. Analysis was carried out using Hi-C matrices binned at 25 kb resolution. Hi-C matrices made from patient and control data mixed in equal proportions were obtained by combining the first halves of each fastq file to create sample “mixA” and the second half of each fastq file to create sample “mixB”. Mixed samples were processed in the same way as the original samples. Insulation scores were calculated following the method of Crane et al. 2015 ^16^ using a window size of eight times the resolution.

Merged datasets from Rao et al. 2014^6^ were downloaded from the 4D Nucleome^17^ website (https://data.4dnucleome.org/) in .mcool format. Hi-C matrices binned at 25 kb resolution were extracted and converted to FAN-C format for downsampling and further analysis. Individual biological replicates were downloaded from the 4D Nucleome^17^ website as pairs files and further processed using FAN-C as described above.

Data from Wutz et al. 2017^7^ was downloaded from the 4D Nucleome website as .mcool files. Hi-C matrices binned at 50 kb resolution were extracted and converted to FAN-C format for downsampling and further analysis.

### CHESS

CHESS version 0.3.7 was run using window sizes of 2 Mb, with a 500 kb step size, for data binned at 25 kb resolution, and 4 Mb windows with a 1 Mb step size for data binned at 50 kb resolution. To reduce computational processing time, only chromosomes 2 and 16 were analysed for the Rao et al. and Wutz et al. datasets.

The implementation of SSIM used by Lee et al.^2^ is slightly different from that in CHESS. Lee et al. calculate SSIM using the mean of SSIM in 7×7 pixel submatrices across a region, while CHESS by default calculates SSIM using the whole region.

### Statistics and visualisation

Statistics were calculated using R version 4.0.3. Visualisation was carried out using ggplot2 ^18^ and FAN-C ^15^.

## Data availability

Data from Diaz et al.^3^ is available from ArrayExpress under accession number E-MTAB-5875. Data from Rao et al.^6^ is available from https://data.4dnucleome.org/ under Experiment Set Accessions 4DNESI7DEJTM (K562) and 4DNES3JX38V5 (GM12878). Data from Wutz et al.^7^ is available from https://data.4dnucleome.org/ under Experiment Set Acessions 4DNES51Q5×3O, 4DNES7QY4JHS, 4DNESIKACYZC, 4DNESJ7ABWFM, 4DNESJAU6DPJ, 4DNESLZVKJ7V, 4DNESR381AXL, 4DNESR8I1SZG, and 4DNESWO4PE7L.

## Code availability

Code is available on Github: https://github.com/vaquerizaslab/chess-2021

## Author contributions

EI-S carried out data analysis. All authors conceptualized the study, interpreted results, and wrote the manuscript.

## Figures and Figure Legends

**Extended Data Figure 1.**
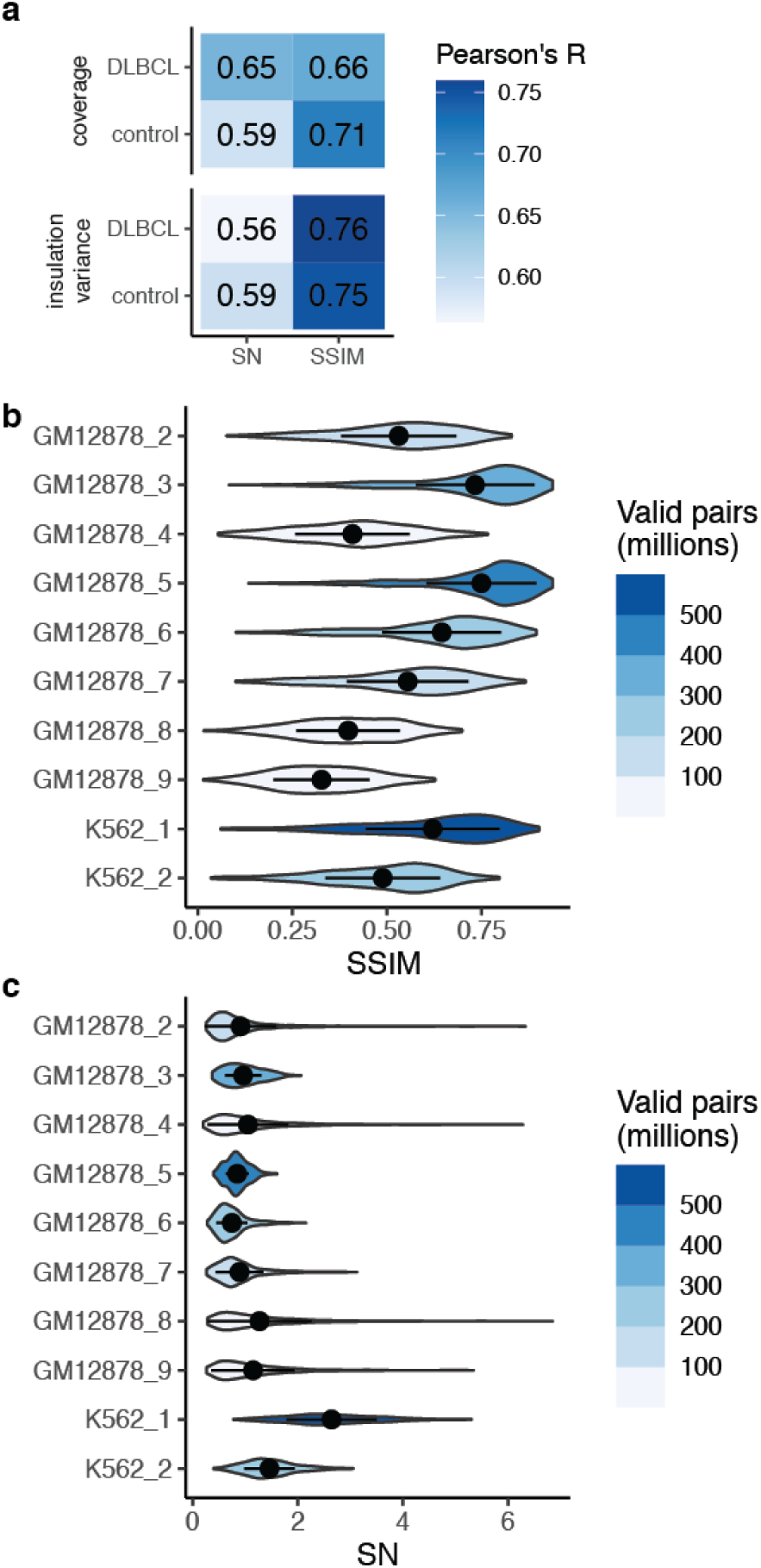
SSIM and SN correlate with genomic coverage and insulation score variance. A. Pearson’s correlations of SSIM and SN profiles from the DLBCL-control comparison with genomic coverage of the Hi-C datasets and the variance of the insulation score in the 2 Mb window used for the CHESS analysis. B. Overall SSIM distributions derived from comparisons of GM12878 and K562 Hi-C data from Rao et al. 2014. In all cases GM12878 biological replicate 1 was used as the reference dataset for the CHESS comparisons. This sample has 1.8 billion valid pairs. Point and lines show mean +/- one standard deviation. C. Overall SN distributions for the same comparisons as in B. Point and lines show mean +/- one standard deviation.

**Extended Data Figure 2.**
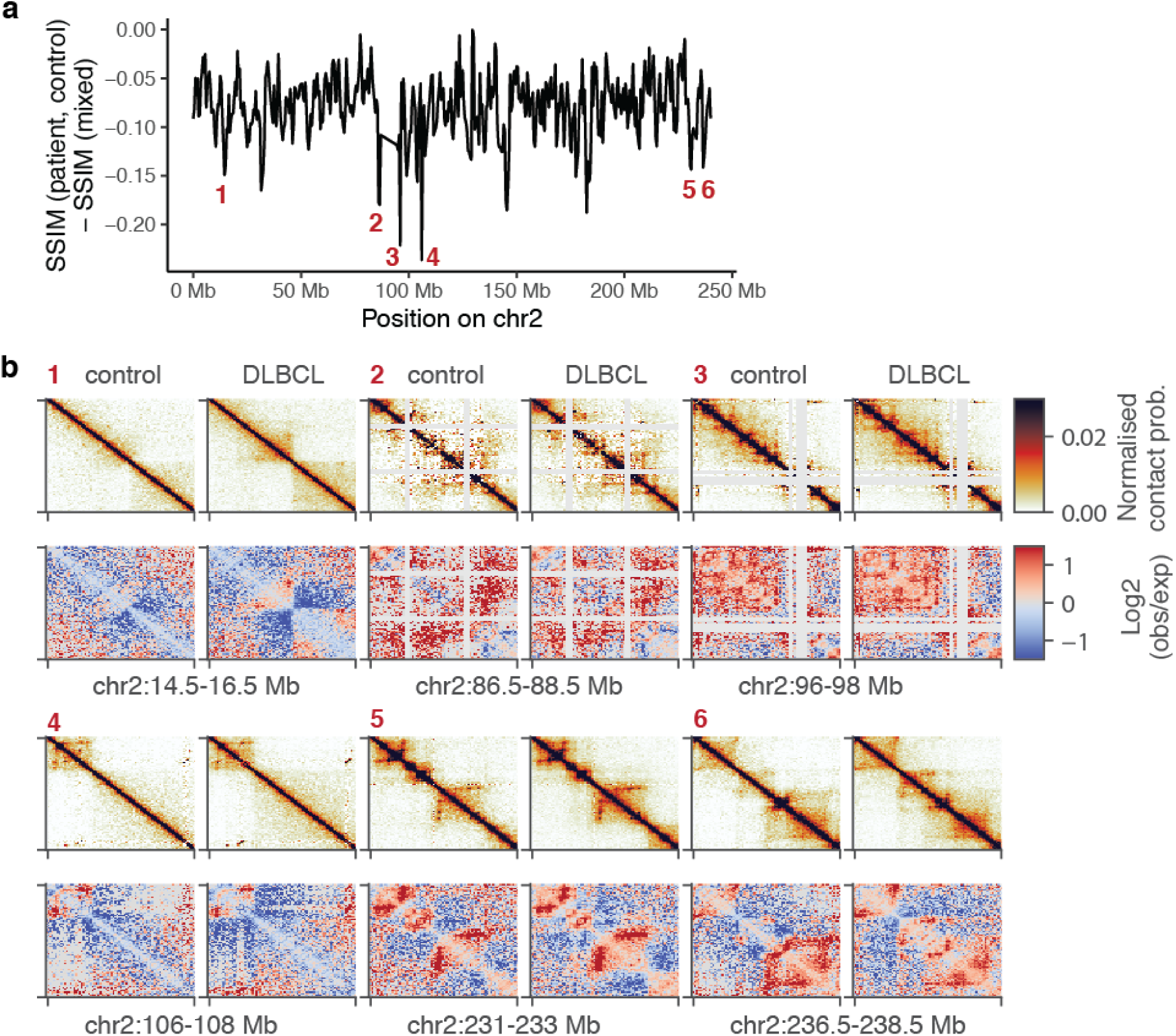
Differences in SSIM between the DLBCL-control comparison and the comparison of mixed datasets highlight changes in genome organisation. A. Subtraction of SSIM (mixed) from SSIM (DLBCL, control) highlights regions with changes between DLBCL and control, reproduced from Figure 1C. B. Examples of regions where the SSIM difference profile indicates differences between DLBCL and control. Top, normalised Hi-C data at 25kb resolution. Bottom, log2 observed / expected values for the same regions. Regions 2 and 3 are adjacent to the poorly-mappable centromeric region of chr2. CHESS performance in these regions may be improved by optimised filtering of low coverage Hi-C bins.

**Extended Data Figure 3.**
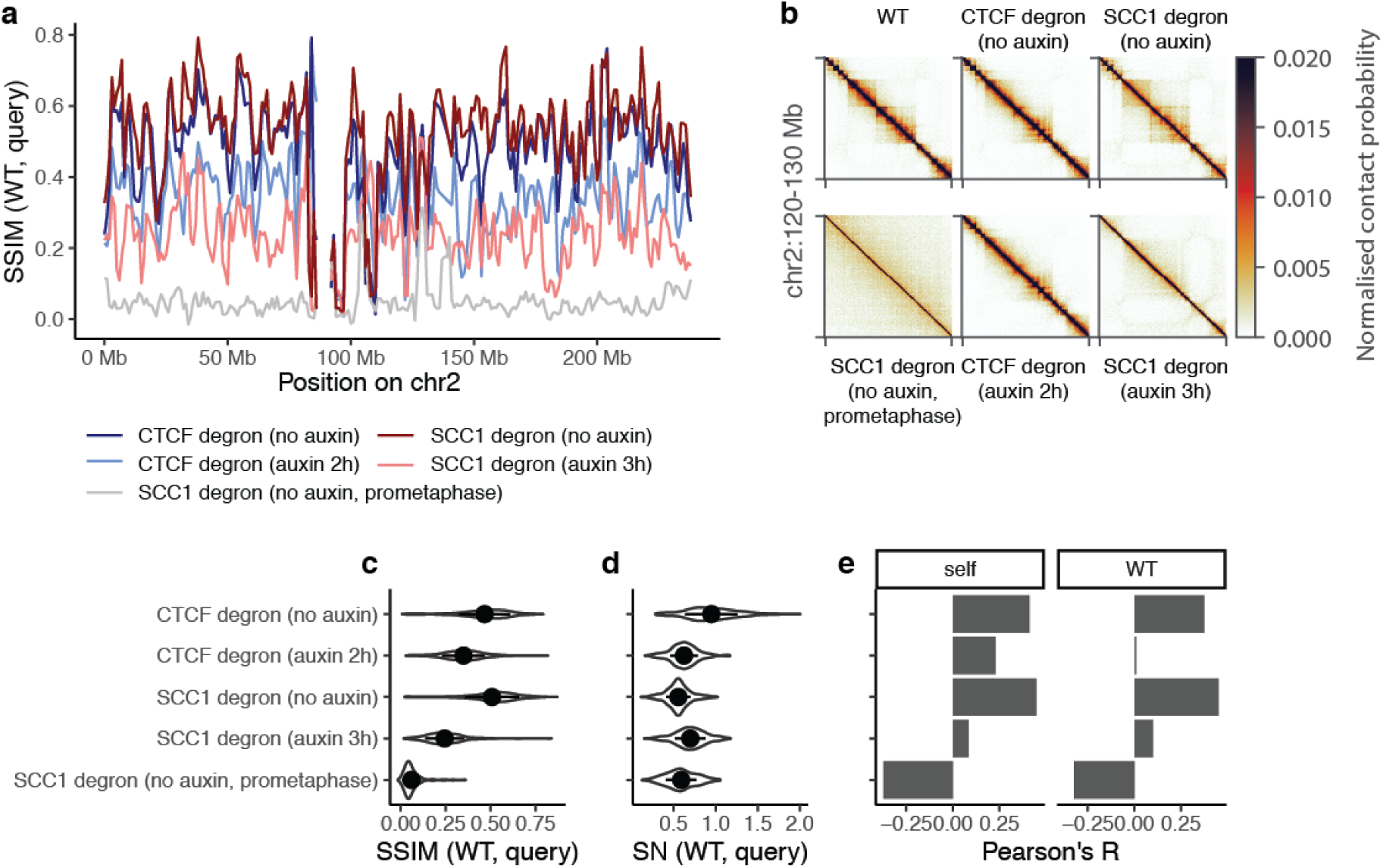
Global changes in SSIM due to global alterations of genome organisation. A. SSIM profiles for comparisons of Hi-C experiments from Wutz et al. 2017^7^. In each case unmodified (“WT”) HeLa cells were used as the reference for the comparison and the query sample is shown in the legend. All cells are synchronised in G1 phase unless otherwise stated. B. Normalised Hi-C data at 50kb resolution for an example region on chr2, for the datasets from A. C. Overall SSIM distributions for the comparisons in A. Point and lines show mean +/- one standard deviation. D. Overall SN distributions for the comparisons in A. Point and lines show mean +/- one standard deviation. E. Pearson’s correlations of the SSIM profile for comparisons in A with genomic coverage of the Hi-C datasets used in the comparison.

